# GAUT10 is required for Arabidopsis root cell differentiation and elongation

**DOI:** 10.1101/2023.02.07.527497

**Authors:** Linkan Dash, Sivakumar Swaminathan, Jan Šimura, Christian Montes, Neel Solanki, Ludvin Mejia, Karin Ljung, Olga A. Zabotina, Dior R. Kelley

## Abstract

- Cell wall properties of the root apical meristem (RAM) are poorly understood compared to the elongation and maturation zones of the developing root. GAUT10 is a pectin biosynthesizing enzyme that is post-transcriptionally regulated by auxin that influences Arabidopsis RAM size in a sucrose-dependent manner.
- Using live-cell microscopy, we have determined the short root phenotype of the *gaut10* loss of function allele is due to a reduction in both RAM cell number and epidermal cell elongation. In addition, the absence of *GAUT10* leads to a reduction in lateral root cap and epidermal cell marker line expression, indicating root cell differentiation defects in this mutant.
- GAUT10 is required for normal pectin and hemicellulose composition in primary Arabidopsis roots. Loss of *GAUT10* leads to a reduction in galacturonic acid and xylose in primary cell walls and alters the presence of rhamnogalacturonan (RG) I and homogalacturonan (HG) polymers in the root.
- Auxin mediated gene expression and metabolism is altered in *gaut10* roots, suggesting that cell wall composition may influence auxin pathways.

## Introduction

Roots are a key organ in plants for water and nutrient acquisition. In angiosperms, the primary root is established during embryogenesis and further elaborated post-embryonically as stem cells differentiate and daughter cells grow. *Arabidopsis* has served as a foundational model for root growth and development, whereby the cells that have established root identity during embryogenesis continue to grow post-embryonically because of well-regulated stem cell activities (Clark *et al*., 2019a). Within the primary root apical meristem, the stem cell niche consists of the mitotically inactive quiescent center (QC) and several types of progenitor cells, including distal columella cells, lateral root cap/epidermal initials (Epi/LRC), cortex/endodermal initials, and vasculature initials. These progenitor cells further differentiate into specific cell types that mature via cell elongation (Fisher & Sozzani, 2016). An outstanding question in the field is how root morphogenesis is coordinately influenced by internal growth cues, such as hormones, and cellular properties such as cell wall composition. While it is well established that pectin composition varies across root development (Somssich *et al*., 2016), our current understanding of which *Arabidopsis* enzyme(s) underpin root apical meristem (RAM) properties is insufficient.

The plant cell wall is a dynamic structure during development and can exhibit distinct properties based on cellular function. In addition, the plant cell wall provides physical strength to the plant cells and can protect them from internal factors like turgor pressure and external factors like pathogens (Ridley *et al*., 2001). Because plant cells are held in place through cell walls, the stem cell niche must monitor the state of the cell wall. While the transcription factor networks regulating the root stem cell populations have been established over time with extensive genetic studies (Sozzani & Iyer-Pascuzzi, 2014; Fisher & Sozzani, 2016), it is essential to understand the mechanisms beyond transcriptional regulation. The transition of root stem cells across the developmental states leading to their maturation involves regulated cell elongation. However, the essential molecular regulators underlying the cell wall remodeling for such changes in cell morphology are not well understood.

In most angiosperms, including eudicots and non-graminaceous monocots, approximately 35% of the primary cell wall is comprised of pectin (Mohnen, 2008). Pectins are a family of polysaccharides, of which ∼70% are covalently linked units of galacturonic acid (GalA) residues (Keegstra *et al*., 1973). Pectins are synthesized in the Golgi apparatus and subsequently transported to the cell wall (Mohnen, 2008), where they regulate cell wall properties such as extensibility and thickness (Majda & Robert, 2018). The three major classes of pectins that are extensively studied are: homogalacturonan (HG), rhamnogalacturonan-I (RG-I), and substituted galacturonans that are further subclassified as rhamnogalacturonan-II (RG-II), xylogalacturonan, and apiogalacturonan (Scheller *et al*., 2007; Mohnen, 2008).

Current research has limited information on the cell wall properties of root meristem cells. For example, the differences in cell wall composition of mitotically dormant QC cells, surrounding stem initials, and the differentiating cell layers (Somssich *et al*., 2016). Differentiating plant cells have a thin primary cell wall that facilitates frequent cell division (Baluska *et al*., 1996). In such dividing cells, callose predominantly constitutes the early stages of the cell plate assembly (Drakakaki, 2015; Miart *et al*., 2014), that is later replaced by a cellulose/hemicellulose network (Drakakaki, 2015). Pectins and hemicellulose further consolidate the cell wall structure by forming a middle lamella that supports intercellular junctions (Drakakaki, 2015). Cells at the transition zone within the primary root have a unique cell wall pectin composition, with an abundance of (1→4)-β-D-galactan that demarks the transition from mitotic activity to cell elongation (McCartney *et al*., 2003).

Galacturonosyltransferases (GAUTs) are a conserved family of enzymes involved in pectin biosynthesis and belong to the Glycosyl Transferase 8 family (GT8) in the CAZy (Carbohydrate-Active Enzymes) database (Cantarel *et al*., 2009). Genetic and biochemical characterization of *gaut* mutants revealed their unique role in modulating pectin and xylan polysaccharide composition in shoot tissues and contributing to shoot phenotypes (Caffall *et al*., 2009; Wang *et al*., 2013; Lund *et al*., 2020; Guo *et al*., 2021; Engle *et al*., 2022). There are fifteen *GAUT* genes annotated in *Arabidopsis* that are phylogenetically classified into seven clades and ten *GAUT-like* (*GATL*) genes (Caffall & Mohnen, 2009). Two *GAUT* members from clade B-2 have been linked to root growth and development, *GAUT10* and *GAUT15*. *GAUT15* is transcriptionally regulated by auxin and is required for root gravitropism (Lewis *et al*., 2013). GAUT10 is regulated by auxin post-transcriptionally and is required for primary root growth and lateral root formation (Pu *et al*., 2019). Loss of *GAUT10* leads to short roots with reduced RAM size in the absence of exogenous sucrose (Pu *et al*., 2019). In addition, GAUT10 has been shown to be localized to the Golgi and is involved in pectin biosynthesis in *Arabidopsis* inflorescence and stem tissues (Caffall *et al*., 2009; Voiniciuc *et al*., 2018; Guo *et al*., 2021). GAUT10 is also important for stomata formation (Guo *et al*., 2021), suggesting that this cell wall modifying enzyme may play numerous roles in plant development.

In this study we have determined the molecular and cellular basis of the *gaut10* short root phenotype using live-cell microscopy, transcriptomics, and auxin metabolite profiling. Loss of *GAUT10* leads to a reduction in cell division in the RAM and impairs epidermal cell elongation in the root. Furthermore, the absence of *GAUT10* impacts auxin-dependent gene expression and metabolism, suggesting that the cell wall composition may influence this hormone pathway indirectly. In addition, we have characterized how GAUT10 contributes to root pectin and hemicellulose composition, which may impact root morphogenesis via both cell elongation and differentiation. This study defines a new role for GAUT10 in root development and expands our understanding of how cell wall properties can influence the RAM.

## Materials and Methods

### Plant Material

All seed stocks used in this study were obtained from the Arabidopsis Biological Resource Center (ABRC) at Ohio State University. *Arabidopsis thaliana* plants used in this study were all in the Columbia (Col-0) background. SALK_092577 corresponds to the *gaut10-3* null allele which has been previously characterized (Pu *et al*., 2019). The *WEREWOLF:GFP* (epidermal cell marker), *702LRC:GFP* (lateral root cap marker), *SCARECROW:GFP* (cortex/endodermal cell marker), *PET111:GFP* (columella cell marker), and *DR5:GFP* lines have been previously described (Friml *et al*., 2003; Heisler *et al*., 2005; Bargmann *et al*., 2013; Hayashi *et al*., 2014).

For phenotyping assays, seeds were surface sterilized using 50% bleach and 0.01% Triton X-100 for 10 minutes and then washed five times with sterile water. Seeds were then imbibed in sterile water for two days at 4°C and then transferred to 0.5X Murashige and Skoog medium (MS) plates supplemented with 15 mM sucrose and 0.8% agar. Seedlings were grown under long-day photoperiods (16 h light/8 h dark) at 23°C in a Percival growth chamber.

### Root phenotyping

Intact seedlings (five-day-old) were imaged on a flat-bed scanner (Epson V600). Measurements of primary root lengths were performed using ImageJ. Roots were stained with propidium iodide (PI) and imaged under a confocal microscope for root meristem and differentiation zone phenotyping using a 20X objective. Measurements of root meristem lengths (apical and basal) and the epidermal cells at the differentiation zone were performed using ImageJ, and the number of cortical cells was counted across six biological replicates as previously described (Hacham *et al*., 2011). Two-sample nonparametric Wilcoxon Rank Sum test were performed to compare each phenotype of *gaut10-3* against control Col-0.

### Genotyping

All primers used for genotyping are provided in Table S6. Primers to genotype SALK alleles were designed using the SALK T-DNA verification primer design tool at the SALK SiGNAL website (http://signal.salk.edu/tdnaprimers.2.html).

### Confocal Microscopy

Homozygous *gaut10-3* plants were crossed with the following transgenic lines: *WEREWOLF:GFP*, *702LRC:GFP*, *SCARECROW:GFP*, *PET111:GFP*, and *DR5:GFP*. After the initial crosses, F1 seedlings were genotyped by polymerase chain reaction (PCR) using T-DNA and transgene specific primers (Table S6) and confirmed to be heterozygous for *GAUT10/gaut10-3* and carrying the correct *GFP* transgenes. F1 plants were selfed and then carried through the F3 generation; genotypes were verified by PCR at each generation. F3 individuals that were homozygous for *gaut10-3* and each GFP reporter were bulked for imaging. Five-day-old roots of these double transgenic and corresponding control transgenic lines were stained with propidium iodide (PI) and imaged under a 20X objective on a Zeiss LSM 700 confocal microscope. 488 and 555 nm lasers were used for GFP and PI excitations, respectively.

### Transcriptomic Profiling

Transcriptomic analyses were performed as previously described (Dash *et al*., 2021). Col-0 and homozygous *gaut10-3* seeds were surface sterilized using 50% bleach and 0.01% Triton X-100 for 10 minutes and then washed five times with sterile water. Seeds were then imbibed in sterile water for two days at 4°C and then transferred to 0.5X MS medium plates supplemented with 15 mM sucrose and 0.8% agar and overlaid with sterile 100-micron nylon mesh squares to facilitate tissue harvesting. Seedlings were grown under long-day photoperiods (16 h light/8 h dark) at 23°C. Roots from five-day-old seedlings were dissected using surgical knives and then weighed and snap-frozen in liquid nitrogen; approximately 100 mg of seedling root tissue was collected per replicate/genotype. Three independent biological replicates were generated for each genotype. Snap frozen roots were ground to a fine powder in liquid nitrogen using a mortar and pestle. Total RNA was extracted from the root samples using Trizol, followed by column clean-up using the Quick-RNA plant kit (Zymo research). Total RNA concentration was estimated using a NanoDrop and Qubit. RNA quality was checked via Bioanalyzer at the ISU DNA Facility. QuantSeq 3’ mRNA libraries were prepared using the Lexogen 3’ mRNA-seq FWD kit and sequenced on an Illumina HiSeq 3000 as 50 bp reads at the ISU DNA Facility. QuantSeq reads were mapped to the TAIR10 genome using STAR (Dobin *et al*., 2013), and differential gene expression analysis was performed using DeSeq2 implemented in R (Love *et al*., 2014). Transcripts with an FDR cutoff of < 0.05 and a log_2_fold change ≥ 0.5 were defined as differentially expressed (DE) (Table S3).

### Gene ontology analysis

Differentially expressed genes (DEGs) were tested for statistical over-representation of Gene Ontology (GO) biological process (GOBP) terms using Panther and the *Arabidopsis thaliana* database. A standard fisher’s exact test with FDR correction was used to calculate GO term enrichment against the total set of detected transcripts as the reference (Table S3). Top GOBP terms were plotted on the y-axis against their genotype/treatments on the x-axis in multidimensional dot plots.

### Accession Numbers

Raw RNA sequence read data from this article can be found at the NCBI BioProject database under Accession: PRJNA694693 and ID: 694693.

### Cell wall, pectin, and hemicellulose extraction

Total cell wall material, pectin, and hemicellulose fractions from three biological replications of Col-0 and *gaut10-3* root samples were prepared as previously described (Zabotina *et al*., 2012). Briefly, the root samples were ground to a fine powder in liquid nitrogen, and the cell wall was extracted by using successive organic solvents, starting with 80% (v/v) ethanol, followed by 80% (v/v) acetone, chloroform: methanol (1:1, v/v) and finally with 100% acetone. The resultant cell wall material was air dried, and the pectin and hemicellulose polysaccharides were extracted successively by using 50 mM cyclohexane diamine tetra acetic acid (CDTA): 50 mM ammonium oxalate (1:1) buffer (v:v) and 4M KOH, respectively. The extracts were neutralized with acetic acid, dialyzed against water, and dried by lyophilization. The dried pectin-enriched and hemicellulose-enriched extracts were stored at 4°C for further analysis.

### Glycome profiling of epitopes of pectin and hemicellulose extracts

Glycome profiling was performed as previously described (Pattathil *et al*., 2010, 2012). The amount of sugar in the pectin and hemicellulose extracts was determined by the phenol-sulfuric acid method, and each enzyme-linked immunosorbent assay (ELISA) well was loaded with an equal amount of polysaccharide sample dissolved in water (50 µl/well from a 60 µg/µL solution). Glycome profiling was carried out using 66 different mouse-primary glycome antibodies purchased from the Complex Carbohydrate Research Center (University of Georgia) as previously described (Pattathil *et al*., 2010, 2012; Zabotina *et al*., 2012). The color development was detected at 450 nm wavelength using a plate reader, and each sample’s OD reading was statistically analyzed and compared. The ELISA assays were repeated in triplicate for both the pectin and hemicellulose extracts. Statistical analysis was done using a non-parametric Wilcoxon rank sum test with a *P* **≤** *0.1* to compare mean OD_450_ absorbances between Col-0 and *gaut10-3*.

### Monosaccharide composition analysis

Monosaccharide composition was determined according to previously described methods (Brenner *et al*., 2012). One mg of cell wall was hydrolyzed with 2M trifluoroacetic acid (TFA) and analyzed by high-performance anion-exchange chromatography with pulsed amperometric detection (HPAEC–PAD) (Dionex, Sunnyvale, CA) using a CarboPac PA20 column. The column was calibrated using monosaccharide standards purchased from Sigma–Aldrich, which included L-Fuc (L-fucose), L-Rha (l-rhamnose), L-Ara (l-arabinose), D-Gal (D-galactose), D-Glc (D-glucose), D-Xyl (D-xylose), D-Man (D-mannose), D-GalA (D-galacturonic acid) and D-GlcA (D-glucuronic acid) (Brenner *et al*., 2012). The content of different monosaccharides in the hydrolysates analyzed was estimated as mol%. Statistical analysis was done using a non-parametric Wilcoxon rank sum test with a *P* **≤** *0.1* to compare mean monosaccharide concentrations between Col-0 and *gaut10-3*.

### Auxin metabolite profiling

Auxin metabolite profiling was performed as previously described (Novák *et al*., 2012). Five-day-old *gaut10-3* and Col-0 roots grown on 0.5X MS + 0.5% (15 mM) sucrose were harvested in pools of 25 mg and flash frozen; 5 biological replicates were analyzed for each genotype. Extraction and liquid chromatography followed by mass spectrometry (LC-MS) was subsequently performed as published. Statistical analysis was done using a non-parametric Wilcoxon rank sum test with a *P* **≤** *0.1* to compare mean auxin metabolite concentrations between Col-0 and *gaut10-3*.

## Results

### GAUT10 is required for root cell division and elongation

Through quantitative proteomics coupled with a reverse genetic screen, we previously identified GAUT10 as an auxin down-regulated protein required for root development (Pu *et al*., 2019; Clark *et al*., 2019b). Loss of function alleles of *gaut10* exhibit a short RAM phenotype compared to wild-type Col-0 when grown in 0.5X MS media lacking sucrose (Pu et al., 2019) and the *gaut10-3* null allele has been well characterized (Caffall *et al*., 2009; Pu *et al*., 2019; Guo *et al*., 2021). Supplementing the 0.5X MS media with 1% sucrose rescues the *gaut10-3* RAM to wild-type size (**Fig. 1**). To determine permissive root growth conditions for analyzing *gaut10-3* roots, we grew Col-0 and *gaut10-3* seedlings on 0.5X MS media supplemented with 0, 0.5%, and 1% sucrose (**Fig. 1**). Based on this assay, we determined that five-day-old *gaut10-3* roots are statistically shorter than Col-0 when grown on 0.5X MS supplemented with 0.5% (15 mM) sucrose, but still long enough to facilitate molecular and biochemical analyses.

**Figure 1.**
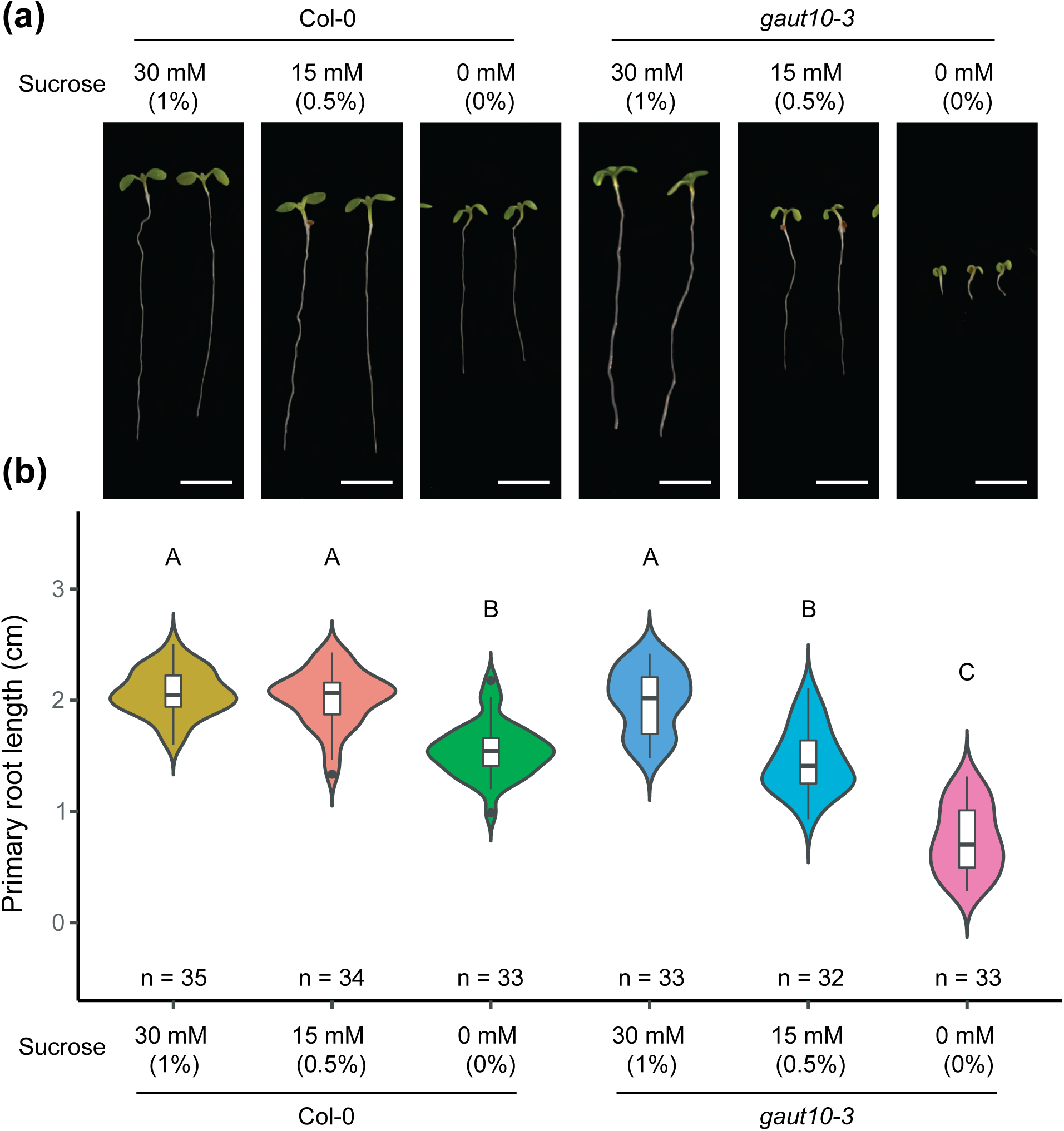
Primary root length is shorter in *gaut10-3* compared to Col-0 grown in 0-and 15-mM sucrose; supplementing the growth media with 30-mM sucrose rescues the root length phenotype. (a) Five-day-old *gaut10-3* and Col-0 whole seedlings grown on 0.5X MS supplemented with three different sucrose concentrations: 0, 15, and 30 mM. Scale bar = 0.5 cm. (b) Violin plots of quantified root phenotypes in Col-0 and *gaut10-*3. Statistical analysis was performed using a one-way ANOVA followed by Tukey’s posthoc analysis with a *P value* < 0.05 to assign letters (A, B, and C) to each genotype/treatment that indicates statistical significance. “n” represents the number of biological replicates quantified. Sucrose concentrations are displayed as both a percentage (weight/volume) and molarity (mM).

Compared to Col-0, *gaut10-3* roots are shorter under 0.5% sucrose conditions (**Fig. 1**). Here, we statistically determined the differences in root lengths via a one-way ANOVA followed by a Tukey’s Post hoc analysis. To determine if this short root phenotype is due to a reduction in cell division and/or elongation, the number of cells in the RAM and the size of mature epidermal cells were quantified as previously described (Hacham *et al*., 2011). Compared to Col-0, the number of apical and basal root meristem cells were reduced in *gaut10-3* (**Fig. 2c-h**). At the transition from the elongation zone (EZ) to the differentiation zone (DZ), *gaut10-3* trichoblast cells are shorter in length compared to wild-type (**Fig. 2a,b,i**). The differences in meristem properties between *gaut10-3* and Col-0 were statistically determined using nonparametric Wilcoxon rank sum tests. Altogether these data suggest that *gaut10* roots are shorter due to both a reduction in cell number and cell elongation.

**Figure 2.**
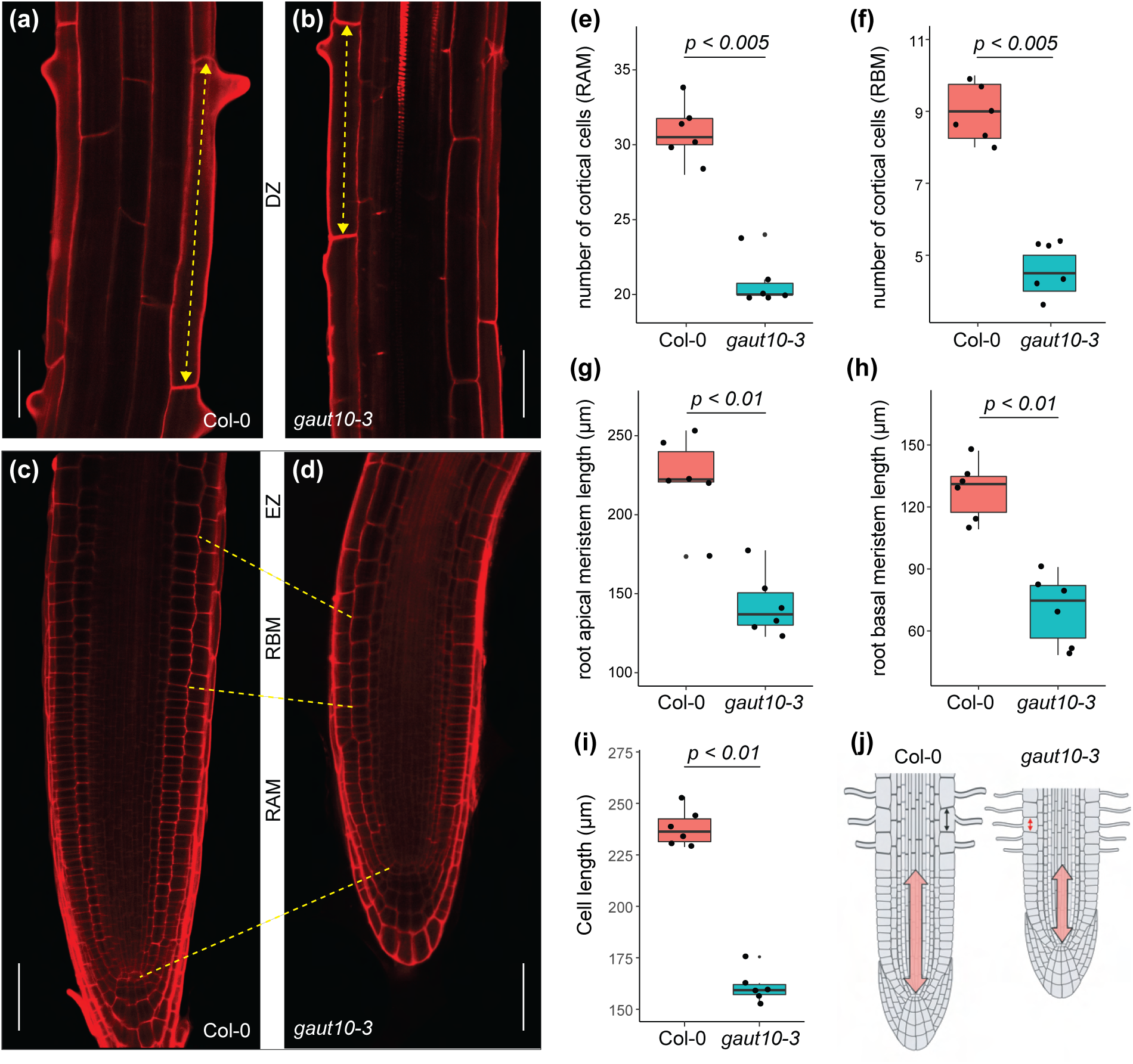
The short phenotype of *gaut10-3* roots is due to a smaller RAM and reduced cell elongation. The confocal images (20X magnification) in panels (a,b) and (c,d) show the root differentiation zone (DZ) and the root meristematic zone (MZ) of five-day-old *gaut10-3* and Col-0 seedlings, respectively, with all scale bars showing 20 μm length; the dotted double head arrows in (a,b) represent the difference in the length of earliest trichoblast cells at the root DZ, the dotted lines in (c,d) demark the length of the root apical and basal meristems in *gaut10-3* and Col-0 representative figures. Phenotypic differences observed in *gaut10-3* as compared to Col-0 are quantified as box and whisker plots overlayed with scatter plots showing differences in the lengths (g,h) and the number of cortical cells (e,f) spanning the apical and the basal root meristems. The meristem lengths are measured on a micrometer (μm) scale. The calculated *P* values are from a non-parametric Wilcoxon Rank-Sum Test.

### Epidermal and lateral root cap marker gene expression is diminished in *gaut10-3*

*GAUT10* mRNA levels have been previously shown to be enriched in several root stem cell types, including the quiescent center, columella, and Epi/LRC (Clark et al., 2019, **Fig. S1**). To determine if loss of *GAUT10* impacts particular cell types withing the RAM, we examined the expression patterns of root marker lines for such cell types. For this experiment, wild-type and *gaut10-3* five-day-old roots harboring root cell marker lines were grown in the absence (-suc) or presence of 0.5% sucrose (+suc) and imaged via confocal microscopy. The QC marker *WOX5:GFP* is expressed normally in *gaut10-3* (**Fig. 3b**), indicating that the stem cell organizing center is normal. However, five-day-old *gaut10-3* roots show diminished levels of *SCR:GFP* in a sucrose-dependent manner, which marks cortex/endodermal cell identity (**Fig. 3a**). In addition, both *PET111:GFP* and *WER:GFP* marker lines showed diminished expression in *gaut10-3* in a sucrose dependent manner (**Fig. 3c,e**), which mark the columella and epidermis respectively. The lateral root cap marker *702LRC:GFP* was absent in *gaut10-3* independent of sucrose (**Fig. 3d**). Altogether, these data suggest that *GAUT10* may influence RAM marker gene expression in a sucrose dependent manner.

**Figure 3.**
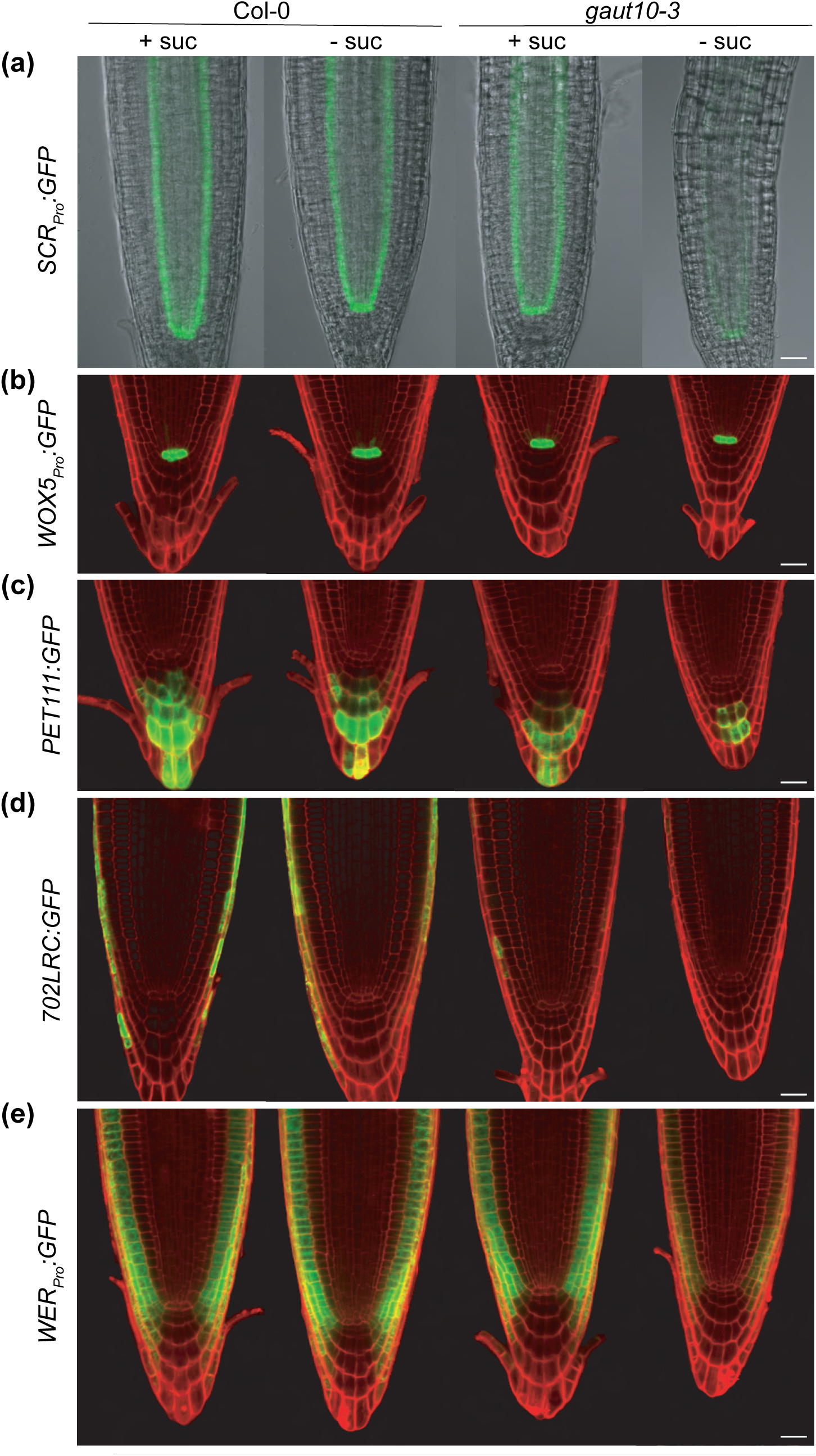
Root cell marker lines are altered in *gaut10*. (A-E) Confocal images of *SCR:GFP* (a)*, WOX5:GFP* (b)*, PET111:GFP* (c)*, 702LRC:GFP* (d)*, WER:GFP* (e) in Col-0 and *gaut10-3* +sucrose (15 mM concentration) and -sucrose. Scale bars = 20 µm.

### Contributions of GAUT10 to cell wall composition

GAUT10 has been previously shown to contribute to the pectin biosynthesis of Arabidopsis shoot tissues (Caffall *et al*., 2009; Guo *et al*., 2021). To determine how the loss of GAUT10 may impact pectin composition in the root, we performed both monosaccharide composition analysis and glycome profiling (**Fig. 4**). The differences in glycosyl residue compositions between *gaut10-3* and Col-0 were statistically determined using a nonparametric Wilcoxon rank sum test. Galacturonic acid (GalA) levels are reduced in the pectin-enriched fraction of *gaut10-3* roots (**Fig. 4a**; Table S2), which is consistent with a previous report on reduced GalA levels of *gaut10* siliques (Caffall *et al*., 2009). In addition, xylose levels are significantly reduced in the hemicellulose-enriched fraction of *gaut10-3* roots (**Fig. 4b**; Table S2).

**Figure 4.**
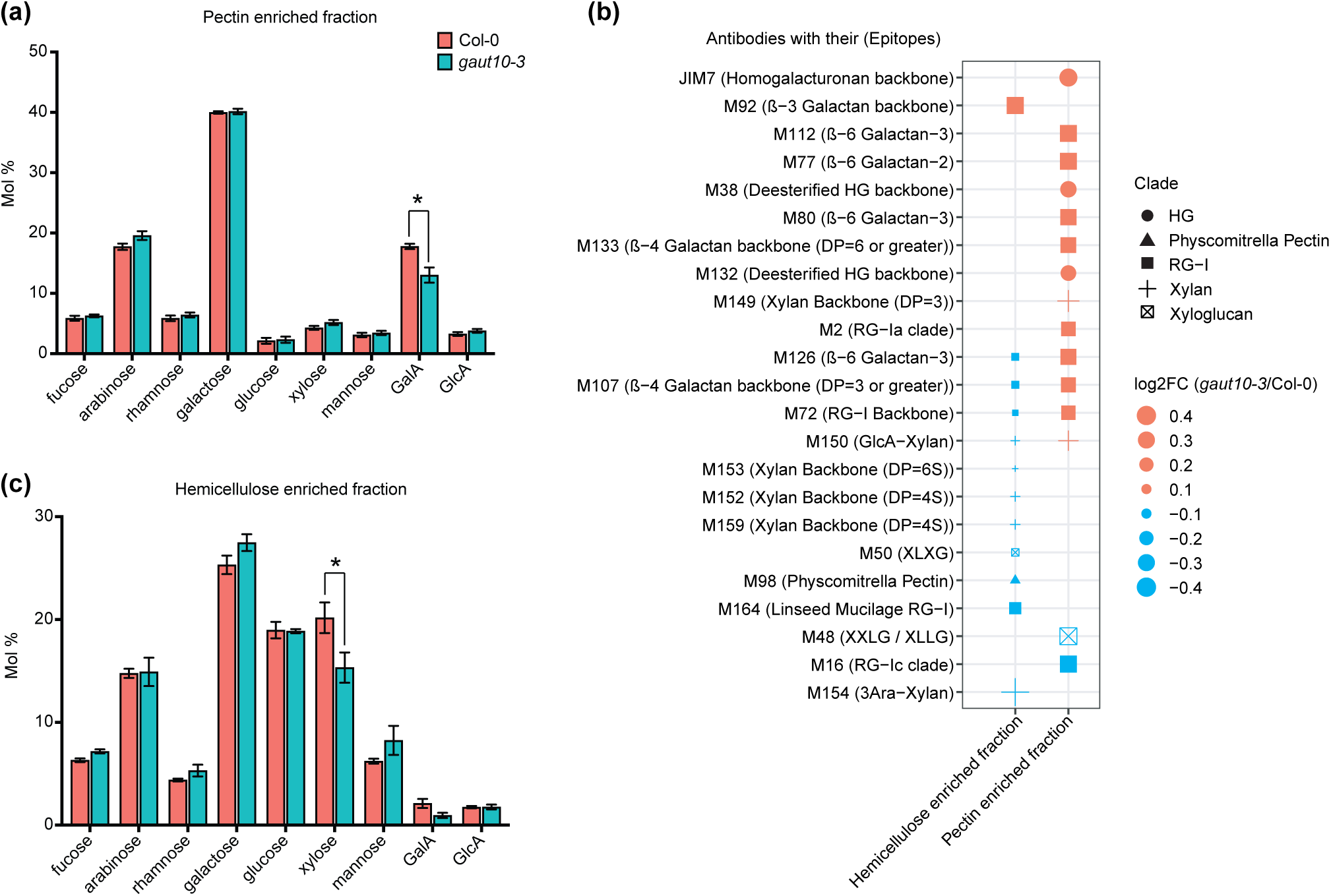
Cell wall polysaccharide composition is altered in *gaut10-3* roots. (a, b) Concentrations of root cell wall monosaccharides (shown as percent molecular weight, “Mol%”) in the pectin (a) and the hemicellulose (b) enriched fractions from three biological replicates per genotype. The error bars show the standard error of the mean (SEM). Statistical analysis was performed using a two-sample nonparametric Wilcoxon rank sum tests; an asterisk indicates *P* value ≤ 0.1. (c) A four-dimensional dot plot that shows differential binding of cell wall polysaccharide-specific antibodies in root cell-wall extracts collected across three biological replicates from *gaut10-3* and Col-0 measured by ELISA. The shape annotations show clades of the cell wall binding antibodies (as defined in Pattathil et al., 2010) whereas the x-axis indicates the corresponding cell wall fractions origin (i.e. pectin or hemicellulose). The size of the dots represents the log2 fold change (*gaut10-3*/Col-0) value calculated using the ELISA absorbance values. Significantly enriched/reduced polysaccharides (23 in total) with their binding epitopes are indicated according to Pattathil et. al., 2010 annotations. Statistical significance is determined by two sample nonparametric Wilcoxon rank sum tests with *P* value ≤ 0.1. Antibody names and their binding cell-wall epitope structures are annotated for each row. Abbreviations used: Homogalacturonan (HG) and Rhamnogalacturonan-I (RG-I).

Glycome profiling was performed using an ELISA based method on the pectin, and the hemicellulose enriched fractions with a large collection of 66 cell wall specific antibodies on 5-day-old Col-0 and *gaut10-3* roots grown on 0.5X MS supplemented with 0.5% sucrose. From these analyses, we identified 15 differentially enriched polysaccharide epitopes in the pectin enriched fraction, and 12 in the hemicellulose enriched fraction of *gaut10-3* root cell walls as compared to Col-0 using statistical comparison of their mean absorbance by a Wilcoxon rank sum test (with *P value* ≤ 0.1) (**Fig. 4c**; Table S2).

The epitopes present in the Homogalacturonan backbone (methyl esterified and de-methyl esterified), Rhamnogalacturonan-Is (RG-I backbone, RG-Ia, RG−I/Galactans), Xylan backbone, and GlcA-Xylan were enriched in the CDTA/Ammonium oxalate soluble pectin abundant fraction from *gaut10-3* root cell walls in comparison with the same fraction prepared from Col-0 cell walls (**Fig. 4c**; Table S2). In contrast, RG-Ic and Xyloglucan (XXLG/XLLG) are reduced in the pectin-enriched fraction of *gaut10-3* compared to Col-0 (**Fig. 4c**; Table S2). On the other hand, 11 out of 12 significant epitopes were found to be reduced in the hemicellulose-enriched fraction of *gaut10-3* root cell-wall compared to Col-0, including RG-Is (RG-I backbone, RG−I/Galactans, Linseed Mucilage RG−I), Xylans (Xylan backbone, GlcA−Xylan, 3Ara−Xylan), and a Xyloglucan (XLXG). Interestingly, only one significant RG-I epitope, namely, ß−3 Galactan backbone, was found to be enriched in the hemicellulose-enriched cell wall fraction of *gaut10-3* roots in comparison to Col-0 (**Fig. 4c**; Table S2). The observed reduction of xylose in the hemicellulose-enriched fraction (**Fig. 4b**) correlates with the reduction of several xylan-related epitopes in glycome profiling of the same fraction (**Fig. 4c**). Altogether these data provide a novel characterization of GAUT10-dependent cell wall composition within Arabidopsis primary roots.

### Loss of *GAUT10* impacts auxin and cell wall pathway gene expression

To determine how gene expression may be impacted in *gaut10-3*, we performed transcriptomics on 5-day-old *gaut10-3* and Col-0 roots grown on 0.5X MS + 0.5% sucrose. Differentially expressed genes were identified using the DESeq2 package implemented in R (Love *et al*., 2014). With an adjusted *P* value of < 0.05, 109 upregulated and 176 downregulated genes were identified in *gaut10-3* roots relative to Col-0 (**Fig. 5a**, Table S3). Among these differentially expressed genes, we identified several key marker genes that are associated with root development and are consistent with the observed defects in *gaut10* roots. For example, *EXTENSIN18* (*EXT18*), which is required for root growth via cell elongation (Choudhary *et al*., 2015) is down regulated in *gaut10-*3 (**Fig. 5b**). In addition, several known auxin pathway genes are altered in *gaut10-3* roots including *YADOKARI 1* (*YDK1*), *NITRILASE 1 (NIT1), MYB DOMAIN PROTEIN 34 (MYB34), CA^2+^-DEPENDENT MODULATOR OF ICR1 (CMI1)* and *INDOLE-3-ACETIC ACID INDUCIBLE 17/AUXIN RESISTANT 3 (IAA17/AXR3)* (**Fig. 5b**). Collectively, *YDK1*, *NIT1* and *MYB34* are important regulators of auxin metabolism (BARTLING *et al*., 1992; Bartling *et al*., 1994; Bartel & Fink, 1994; Takase *et al*., 2004; Celenza *et al*., 2005; González-Lamothe *et al*., 2012; Lehmann *et al*., 2017). *CMI1* is an auxin-regulated gene that modulates auxin responses in the root meristem via a Ca2+-dependent pathway (Hazak *et al*., 2019). Loss of function of *cmi1* results in a shorter primary root with reduced root meristem size (Hazak *et al*., 2019), which is consistent with the *gaut10* root phenotype. *IAA17/AXR3* encodes for an Aux/IAA transcription factor that represses auxin-inducible gene expression (Nagpal *et al*., 2000; Nakamura *et al*., 2006; Muto *et al*., 2007). Altogether the transcriptomic analysis suggests that the altered cell wall composition in *gaut10* may influence auxin pathway gene expression.

**Figure 5.**
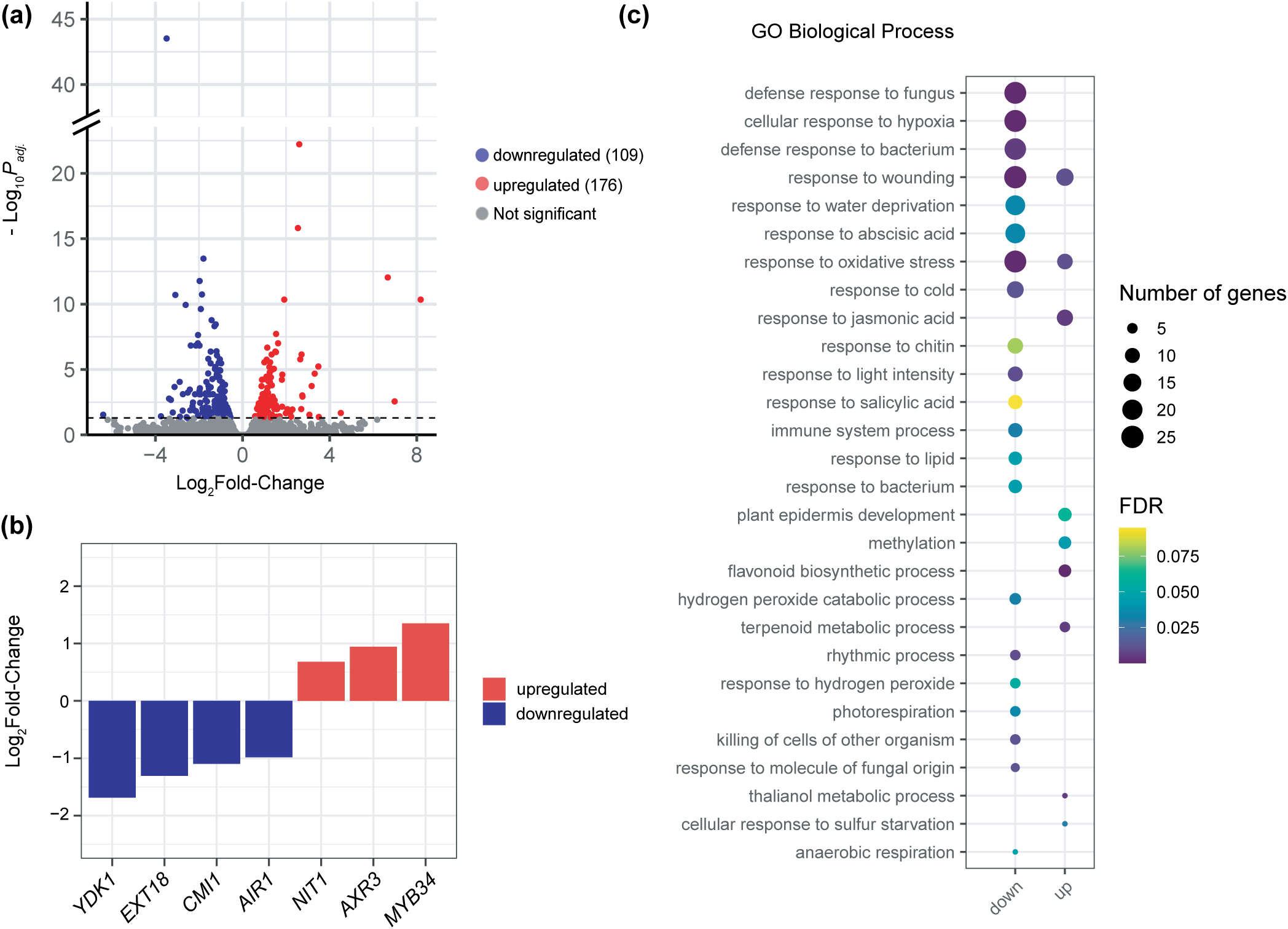
Transcriptomic analysis of *gaut10* primary roots. (a) Volcano plot showing differentially expressed genes with an adjusted *P.adj.* value (FDR) threshold of 0.05, with log_2_Fold-change on the x-axis and -log_10_(adjusted *P.adj. value*) on the y-axis. The upregulated genes (109) are shown in blue, downregulated (176) in red, and non-significant in grey. (b) Expression values for notable key marker genes are shown in the bar plot with log2 fold-change on the y-axis and gene abbreviations on the x-axis: *YADOKARI 1 (YDK1), EXTENSIN 18 (EXT18), CA2+-DEPENDENT MODULATOR OF ICR1 (CMI1), AUXIN-INDUCED IN ROOT CULTURES 1 (AIR1), NITRILASE 1 (NIT1), AUXIN RESISTANT 3 (AXR3), MYB DOMAIN PROTEIN 34 (MYB34).* (c) The top Gene-Ontology (GO) terms for biological processes enriched by the differentially expressed transcripts in *gaut10-3* compared to Col-0 are shown in a four-dimensional graph, where the size and color of the puncta show the number of differentially expressed genes/proteins enriched within each GO term and their statistical significance, respectively.

In order to identify enriched biological processes among DEGs in *gaut10-3* compared to wild-type we performed Gene Ontology (GO) enrichment analysis. Overall, numerous GO terms that are typically associated with plant stress and/or defense are significantly enriched among DEGs in *gaut10-3*. Among upregulated genes there is an enrichment for ‘response to hydrogen peroxide’, and numerous biotic and abiotic stress terms (**Fig. 5c**). In addition, two classical defense and stress hormone GO terms are enriched (‘response to salicylic acid’ and ‘response to abscisic acid’) among the down-regulated genes in *gaut10* (**Fig. 5c**). In contrast, among the up-regulated genes in *gaut10-3* are GOBP terms associated with ‘plant epidermis development’, ‘response to jasmonic acid’ and ‘flavonoid biosynthetic development’ (**Fig. 5c**). This GO analysis suggests that loss of *GAUT10* may impact plant growth and stress responses.

### Local auxin signaling is inhibited in the *gaut10-3* root meristem

The phytohormone auxin regulates much critical growth and developmental cues in plants. Because GAUT10 abundance is regulated by auxin (Pu et. al., 2019) and several auxin pathway genes are DE in *gaut10* roots (**Fig. 5**), including *IAA17/AXR3*, we wanted to examine auxin signaling using the *DR5:GFP* reporter (Friml *et al*., 2003; Heisler *et al*., 2005; Hayashi *et al*., 2014). We observed a significant reduction of *DR5:GFP* expression in *gaut10-3* roots compared to Col-0 (**Fig. 6**), which was exacerbated under sucrose deficit growth conditions (**Fig. 6**; Table S4). Altogether these results suggested a defect in local auxin signaling in the *gaut10-3* root meristem which may be due to the enrichment of *IAA17/AXR3* transcript-a classical repressor for auxin signaling that regulates root architecture (Rouse *et al*., 1998; Nagpal *et al*., 2000; Ouellet *et al*., 2001; Overvoorde *et al*., 2005; Dreher *et al*., 2006; Tian *et al*., 2014; Korasick *et al*., 2014; Yin *et al*., 2022).

**Figure 6.**
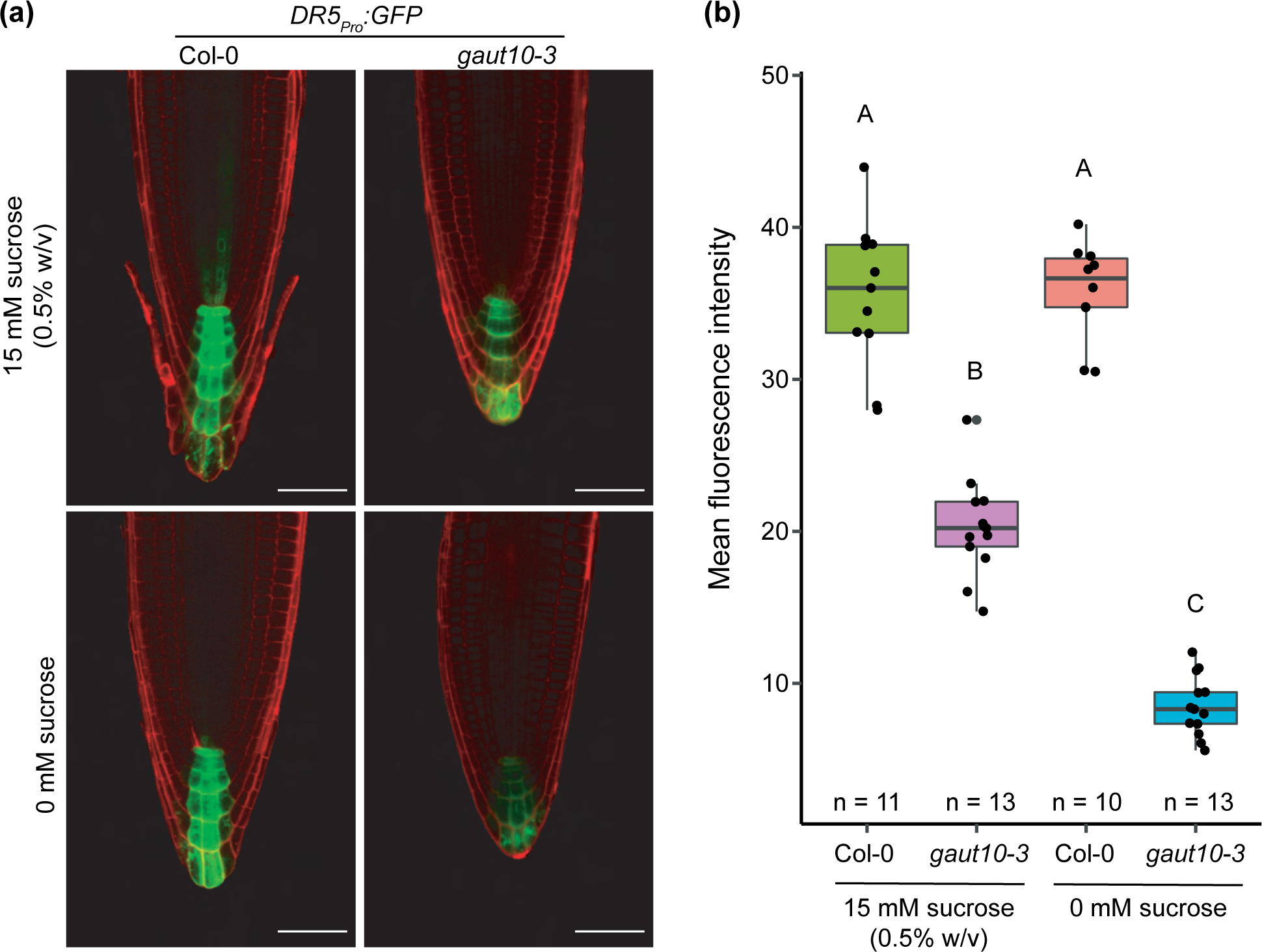
Auxin signaling defects in *gaut10-3* roots compared to wild-type. (a) Representative 20X confocal images of *DR5:GFP* expression in five-day-old Col-0 and *gaut10-3* roots grown on 0 and 15 mM sucrose. Scale bars = 20 µm. (b) Mean fluorescence intensity (MFI) quantified for each genotype/treatment in y-axis. Statistical analysis was a one-way ANOVA followed by Tukey’s posthoc analysis with a *P* value < 0.05 to assign letters (A, B, and C) to each genotype/treatment to indicate statistical significance. “n” represents the number of biological replicates used for averaging.

### Auxin metabolism is altered in *gaut10-3* roots

Auxin homeostasis plays a major role in establishing postembryonic root architecture (Jones & Ljung, 2012). In concert with polar auxin transport, local auxin biosynthesis forms the spatiotemporal auxin gradient within the root apex that plays an essential role in root development (Ljung *et al*., 2005; Petersson *et al*., 2009; Casanova-Sáez *et al*., 2021). Plants maintain auxin homeostasis through *de novo* auxin biosynthesis or by controlling the levels of active auxin through conjugation (primarily amino acids and simple sugars) or by degradation (Normanly, 2010; Ruiz Rosquete *et al*., 2012; Casanova-Sáez *et al*., 2021). We identified several known auxin metabolite genes as DE in *gaut10-3* roots (**Fig. 5b**), suggesting that auxin metabolism and/or signaling may be altered in this mutant. Specifically, *YDK1* conjugates aspartate and other amino acids to IAA, consequently lowering active auxin levels *in vivo* in a tissue-specific manner (Takase *et al*., 2004; González-Lamothe *et al*., 2012). *NIT1* encodes an enzyme that catalyzes the conversion of indole-3-acetonitrile (IAN) to indole-3-acetamide (IAM) as it modulates IAA biosynthesis (BARTLING *et al*., 1992; Bartling *et al*., 1994; Bartel & Fink, 1994; Lehmann *et al*., 2017). *NIT1* overexpression results in shorter primary roots in seedlings, explaining its role in regulating root development via auxin homeostasis (Lehmann *et al*., 2017). In hindsight, a recent review points out the prevailing lack of biochemical and genetic evidence that can attribute these nitrilases (NITs) to indole-3-acetaldoxime (IAOx)-dependent auxin biosynthesis (Casanova-Sáez *et al*., 2021). *MYB34* plays a key role in the transcriptional regulation of enzymes that maintain the metabolic homeostasis of tryptophan and indole glucosinolates in *Arabidopsis* (Celenza *et al*., 2005). To determine how the loss of *GAUT10* may impact auxin metabolism, we measured auxin metabolites in Col-0 and *gaut10-3* roots grown in the presence of 0.5% sucrose using established protocols using LC-MS/MS.

Indole-3-acetic acid (IAA) levels are the same in *gaut10-3* compared to wild-type (**Fig. S3**), but several IAA precursors and conjugates were altered in abundance (**Fig. 7**). The central precursor for auxin biosynthesis in plants-Tryptophan (TRP), that forms indole-3-acetic acid (IAA) via a well-established indole-3-pyruvic acid (IPyA) pathway (Stepanova *et al*., 2008; Tao *et al*., 2008; Yamada *et al*., 2009; Casanova-Sáez *et al*., 2021), was significantly reduced in *gaut10* roots (**Fig. 7b**). Interestingly, Anthranilate (ANT)-the primary rate limiting precursor of TRP biosynthetic pathway was enriched in the mutant roots (**Fig. 7a**). Indole-3-acetonitrile (IAN) that is debatably involved in IAOx to IAA conversion (Sugawara *et al*., 2009; Casanova-Sáez *et al*., 2021), was depleted in *gaut10* roots (**Fig. 7c**). In addition, several inactive forms of IAA were significantly decreased in *gaut10-*3 compared to wild-type, including IAA-Aspartate (IAA-Asp), IAA-Glutamate (IAA-Glu), and IAA-Glucose (IAA-Glc), and 2-oxindole-3-acetic acid-Glucose (oxIAA-Glc) (**Fig. 7d-g**). The observed reduction of IAN and IAA-Asp and increase in ANT is consistent with the differential expression of their respective metabolic enzymes (*YDK1, NIT1*, and *MYB34*) in *gaut10* roots. Collectively, these data suggest that cell wall composition may feedback to alter auxin metabolism within the RAM.

**Figure 7.**
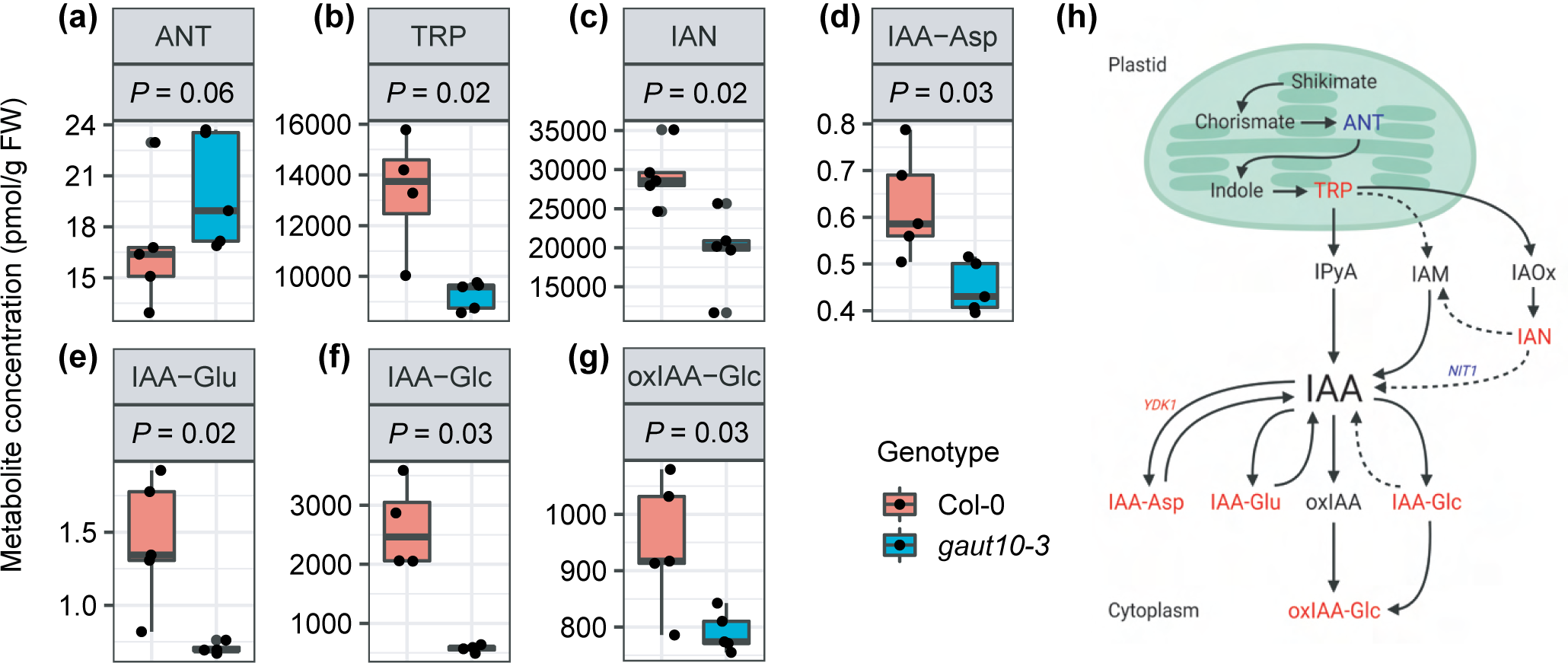
Auxin metabolism is altered in *gaut10-3* roots compared to wild-type. Metabolite quantification by LC-MS/MS visualized by box and whisker plots for with the corresponding P value (*P*) indicated for each metabolite across five biological replicates per genotype. Two sample nonparametric Wilcoxon rank sum tests were done to identify significantly altered metabolites with *P* ≤ 0.1. (a) Anthranilate (ANT) (b) Tryptohan (TRP) (c) indole-3-acetonitrile (IAN) (d) indole-3-acetyl-Aspartate (IAA-Asp) (e) IAA-Glutamate (IAA-Glu) (f) IAA-Glucose (IAA-Glc) (g) oxidized IAA-Glucose (oxIAA-Glc) (h) summary model figure showing annotated auxin biosynthesis pathways. Metabolites in red are reduced in *gaut10-3* roots; likewise, pathway enzymes shown in red are downregulated, whereas the ones in blue show increased transcript levels. The solid indicate pathways that have known enzymes, genes and intermediates, while dashed arrows indicate pathways that are not well defined.

## Discussion

In the elongation zone of the primary root, cells transition from the meristematic zone and rapidly elongate along the longitudinal axis while preventing any lateral expansion of cells that require cell wall remodeling (Ledbetter & Porter, 1963; Schiefelbein & Benfey, 1991; Somssich *et al*., 2016). Cell wall remodeling enzymes largely contribute to such unidimensional cell expansions in response to internal turgor pressure (Ledbetter & Porter, 1963; Cosgrove, 2005). GalA is one of the main monosaccharides constituting all pectin polysaccharides such as HG, RG-I, and RG-II and lesser abandoned xylogalacturonan (Caffall & Mohnen, 2009). Many of the characterized *GAUT* mutants display significant reductions in the amount of GalA present in their cell walls prepared from various above-the-ground parts of the plants (Caffall *et al*., 2009; Voiniciuc *et al*., 2018; Guo *et al*., 2021); however, some mutants like *gaut13* and *gaut14* showed higher content of GalA in their siliques and inflorescences (Caffall *et al*., 2009). To our knowledge, roots have not been analyzed in any *gaut* mutant plants. Here we demonstrated that *gaut10-3* roots also have cell walls with reduced GalA and Xylose. Our results are consistent with previous results obtained for *gaut10-1* and *gaut10-2* inflorescence and siliques (Caffall *et al*., 2009). Double *gaut10 gaut11* plants also showed reduced pectin biosynthesis in stomata cells (Guo *et al*., 2021). Recombinant GAUT10 showed catalytic activity when given UDP-GalA as a substrate; however, the activity was very low, and the authors of the study proposed that GAUT10 may have a unique role in synthesizing either HG or possibly RG polysaccharides compared to other GAUTs (such as GAUT1) (Engle *et al*., 2022).

Monosaccharides such as GalA and Xylose are significantly reduced in the pectin and hemicellulose-enriched cell-wall fractions of *gaut10-3* roots, respectively, which shows a net decrease in GalA and Xylose containing polysaccharides in the *gaut10-3* root cell wall as compared to Col-0. Our glycome analysis of both cell wall fractions observed significant alterations in the levels of specific polysaccharides, which is consistent with our monosaccharide analysis. Antibodies specific to GalA containing RG-I and HG epitopes are enriched in the pectin fraction. We also observed a significant decrease in RG-I, Xylan and Xyloglucan – related epitopes in the hemicellulose-enriched fraction. The increased HG and multiple RG-I pectin-related epitopes, specifically in pectin fraction, might indicate the higher extractability of polysaccharides carrying such epitopes due to their weaker binding nature in the *gaut10-3* cell wall in comparison with cell walls of Col-0. For example, epitopes recognized by four cell wall antibodies (M126, M107, M72 and M150) are increased in the more soluble fraction extracted with CDTA-Ammonium oxalate but consequently reduced in the following extract with strong alkali, suggesting the increased solubility of the corresponding polysaccharide molecules within *gaut10* roots. It is also possible that cell wall reinforcement in *gaut10* roots is loosened, resulting in reduced cell expansion and suppressed growth that is necessary to compensate for the changes in how specific polysaccharides are integrated into cell walls. In the future, more detailed studies will be needed to clarify the impact of loss of GAUT10 on (1) the specific polysaccharide structure/interactions and (2) the general impact on cell wall organization in root cells.

Interestingly, *gaut10* siliques and inflorescences did not show a reduction of xylose (Caffall *et al*., 2009); however, a reduction of both GalA and xylose was observed earlier in another *gaut* mutant-*quasimodo 1 (qua1/gaut8)* (Orfila *et al*., 2005). Pectic galactans have been associated with wall strengthening (McCartney *et al*., 2000), and the higher levels of pectin-specific epitopes in *gaut10-3* mutant roots in comparison with Col-0 roots may also be a result of cell wall structural changes leading to increased cell wall rigidity in mutant root cells resulting in suppression of their growth. Altogether, we can infer that GAUT10 plays a critical role in synthesizing/maintaining these polysaccharides, primarily the pectic polysaccharides within the root cell wall that impacts the organization of cell walls via loosening polysaccharide interactions in the *gaut10-3* (**Fig. 8**).

**Figure 8.**
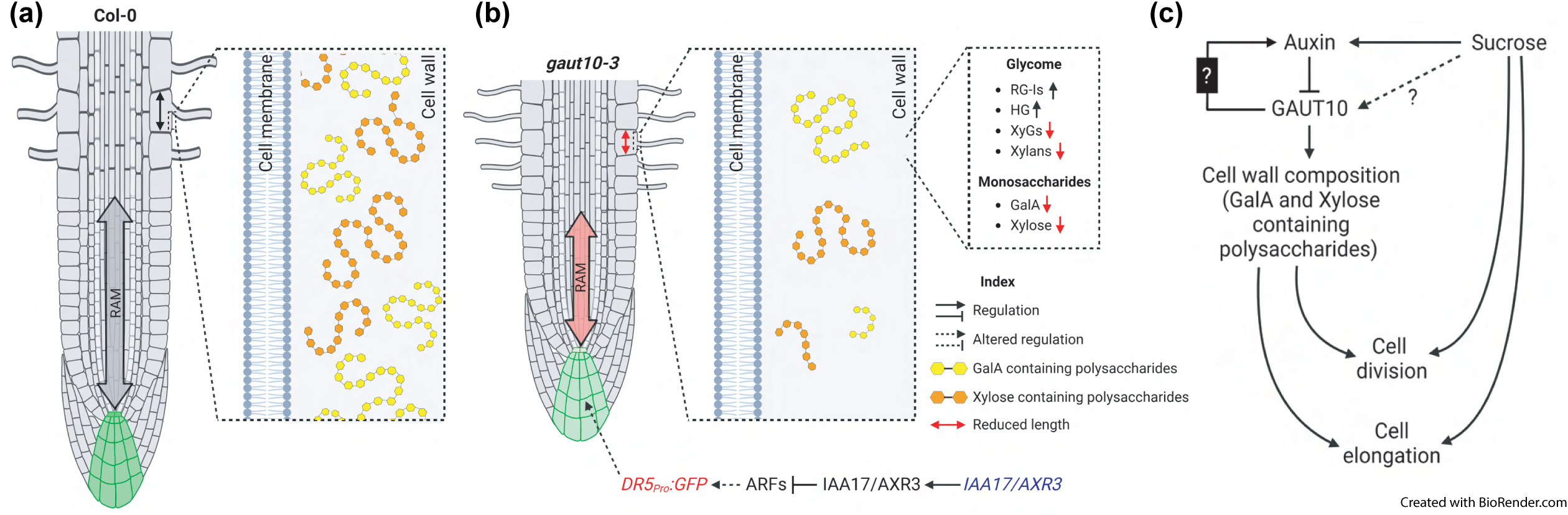
Working model for roles of GAUT10 in root morphogenesis. (a) In wild-type Col-0 roots, normal cell wall composition leads to correct RAM size and DR5:GFP maxima in the QC and columella. (b) In the absence of GAUT10, the cell wall composition is altered leading to reduced cell expansion and root growth. In addition, the expression of *AuxIAA17/AXR3* and *DR5:GFP* are reduced. (c) A molecular framework for GAUT10 function based on biochemical and genetic analyses of *gaut10* roots. Where auxin levels are high in the root, GAUT10 abundance is diminished and cell wall expansion is repressed, such as in the RAM. As cells transition from the RAM to the elongation zone, increased GAUT10 activity promotes cell expansion. Changes in cell wall composition impact auxin metabolism via an unknown mechanism, as indicated by a black box with a question mark. Sucrose is a positive regulator of root morphogenesis via auxin (previous studies) and GAUT10, based on the genetic data provided in this study; the dashed line represents an unknown molecular mechanism.

GAUT10 was identified as an auxin down-regulated protein (Pu *et al*., 2019; Clark *et al*., 2019b) with roles in regulation of RAM size by a reverse genetic study with three loss of function alleles (Pu *et al*., 2019). Here, we performed transcriptomics and metabolomics on *gaut10-3* to substantiate its role as a cell wall enzyme that interacts with auxin pathways and further delineate how GAUT10 function may be linked to auxin biology. We used physiological growth conditions (0.5X MS with 15 mM sucrose) to enable phenotyping and biochemical analysis of *gaut10* roots. The short root phenotype in *gaut10-3* manifest as a combination of smaller root apical meristem with a fewer number of cells and reduced elongation of cells at the differentiation zone, suggesting altered cell division in the meristem as well as precocious maturation of the meristematic cells into the differentiation zone (**Fig. 8c**). The latter conclusion is further supported by overexpression of genes enriching GO term for plant epidermis development in the *gaut10-3* roots.

Root morphogenesis can be studied by visualizing the expression of cell-type specific markers (Bargmann *et al*., 2013). Here, we observed reduced expression of several root cell marker lines (*SCR:GFP*, *WER:GFP*, *PET111:GFP*, and *702LRC:GFP*) in *gaut10* in the absence of sucrose while expression of the QC marker *WOX5:GFP* was normal. Given that numerous plant growth and stress genes are differentially expressed in *gaut10* roots (**Fig. 5**), these data suggest that proper root morphogenesis involves GAUT10-dependent cell wall formation. Future investigations of cell wall composition across the developing primary root may be informative for identifying which cell type specific contexts are associated with particular HG and/or RG moieties.

Cytokinin promotes the degradation of free auxin (IAA) into an inactive conjugate, IAA-Glu, in the lateral root cap, thereby controlling root meristem size and root growth by inducing cell differentiation (di Mambro *et al*., 2019). The diminished expression of *702LRC:GFP* in *gaut10-3* roots that indicates defects in the lateral root cap (**Fig. 3**) might explain the decrease in IAA-Glu (along with other IAA-conjugates) (**Fig. 7**). In addition, modulation of IAA metabolic inactivation and GH3 activity influences root morphogenesis (Mateo-Bonmatí *et al*., 202; Casanova-Sáez *et al*., 2022) . We hypothesize that under physiologically relevant sucrose conditions (0.5% or 15 mM) the growth of *gaut10* roots is restored compared to no sucrose conditions via a reduction in auxin inactivation via conjugation (**Fig. 8**). The observed down regulation of both IAA amino acid (IAA-Asp and IAA-Glu) and sugar conjugates (IAA-Glc), as well as the oxidized forms of IAA (oxIAA-Glc) in *gaut10* roots suggest that IAA homeostasis is restored by down-regulating these metabolic pathways. The differential expression of enzymes catalyzing the conversion of these auxin conjugates and precursors in gaut10 compared to wild-type roots further supports this working model.

Local biosynthesis of auxin in the quiescent center (QC) is sufficient for generating auxin maxima essential for the maintenance of root apical meristem and promoting root growth in *Arabidopsis* (Ljung *et al*., 2005; Brumos *et al*., 2018; Morffy & Strader, 2018). The expression of *DR5:GFP* in *gaut10-3* roots independent of sucrose suggests an auxin signaling defect (**Fig. 6**), further confirmed by the downregulation of auxin-induced genes such as *CMI1, AIR1*, and upregulation of auxin-repressed *AXR3/IAA17* gene in the mutant roots grown in 0.5% sucrose (**Fig. 5b**). These data lead to a working model for GAUT10 which includes auxin signaling and metabolism (**Fig. 8**). Previous studies have shown that along with polar auxin transport, local biosynthesis of auxin plays a critical role in maintaining auxin homeostasis in the root meristem (Ljung *et al*., 2005; Brumos *et al*., 2018). Here, we hypothesize that the cell wall may indirectly impact auxin metabolism via GAUT10 activity (**Fig. 8b, c**).

Sucrose positively regulates root stem cell activation and promotes root epidermal cell elongation in *Arabidopsis* (Dong et al., 2022; Wu et al., 2019; Osorio-Navarro et al., 2022). The sucrose-dependent root growth defects of *gaut10* suggest a potential link between cell wall composition and sucrose-dependent growth that could be explored in future studies. In addition, the downregulation of known sugar transporter genes in *gaut10-3* roots (Table S5) supports an alternative hypothesis whereby cell wall composition may impact apoplastic sugar transport. In summary, GAUT10 is an auxin-downregulated protein that plays a critical role in the synthesis/maintenance of GalA- and Xylose-containing cell wall polysaccharides in the *Arabidopsis* roots, thereby influencing cell wall composition that is required for proper auxin metabolism and morphogenesis within the primary root (**Fig. 8**).

## Data availability

The transcriptome data that support the findings of this study are openly available at the NCBI BioProject database under Accession: PRJNA694693 and ID: 694693. The phenotyping, glycome, and auxin metabolite data are available within the supplementary materials. All seed stocks are available upon request.

## Acknowledgments

This study was supported by NSF Award Number 2118253 to D.R.K. and O.A.Z. We wish to thank Bastiaan Bargmann for sharing seed stocks. KL and JŠ were funded by the Swedish Research Council (VR 2018-04235), the Knut and Alice Wallenberg Foundation (KAW 2016.0341 and KAW 2016.0352) and the Swedish Governmental Agency for Innovation Systems (VINNOVA 2016-00504). We also acknowledge the Swedish Metabolomics Centre for technical support.

## Conflict of interest

We have no conflicts of interest to disclose.

## Author contributions

L.D., S.S., J.S., K.L, O.A.Z, and D.R.K. were involved in design of the research. L.D., S.S., J.S., C.M., N.S., and L.M. performed the research. Data analysis, collection and interpretation was performed by L.D., S.S., J.S., O.A.Z., and D.R.K. The manuscript was written by L.D., S.S., O.A.Z. and D.R.K. with input from the other authors.

Table S1. Root phenotyping data of five-day-old gaut10-3 and Col-0 roots.

Table S2. Glycome and Monosaccharide composition data of five-day-old gaut10-3 and Col-0 roots.

Table S3. Transcriptomic analysis data from five-day-old *gaut10-3* and Col-0 roots with gene expressions averaged across three biological replicates.

Table S4. DR5:GFP fluorescence intensity data.

Table S5. Auxin metabolite concentrations of five-day-old *gaut10-3* and Col-0 roots averaged across five biological replicates.

Table S6. Primers used in this study.

---

**Figure S1.**
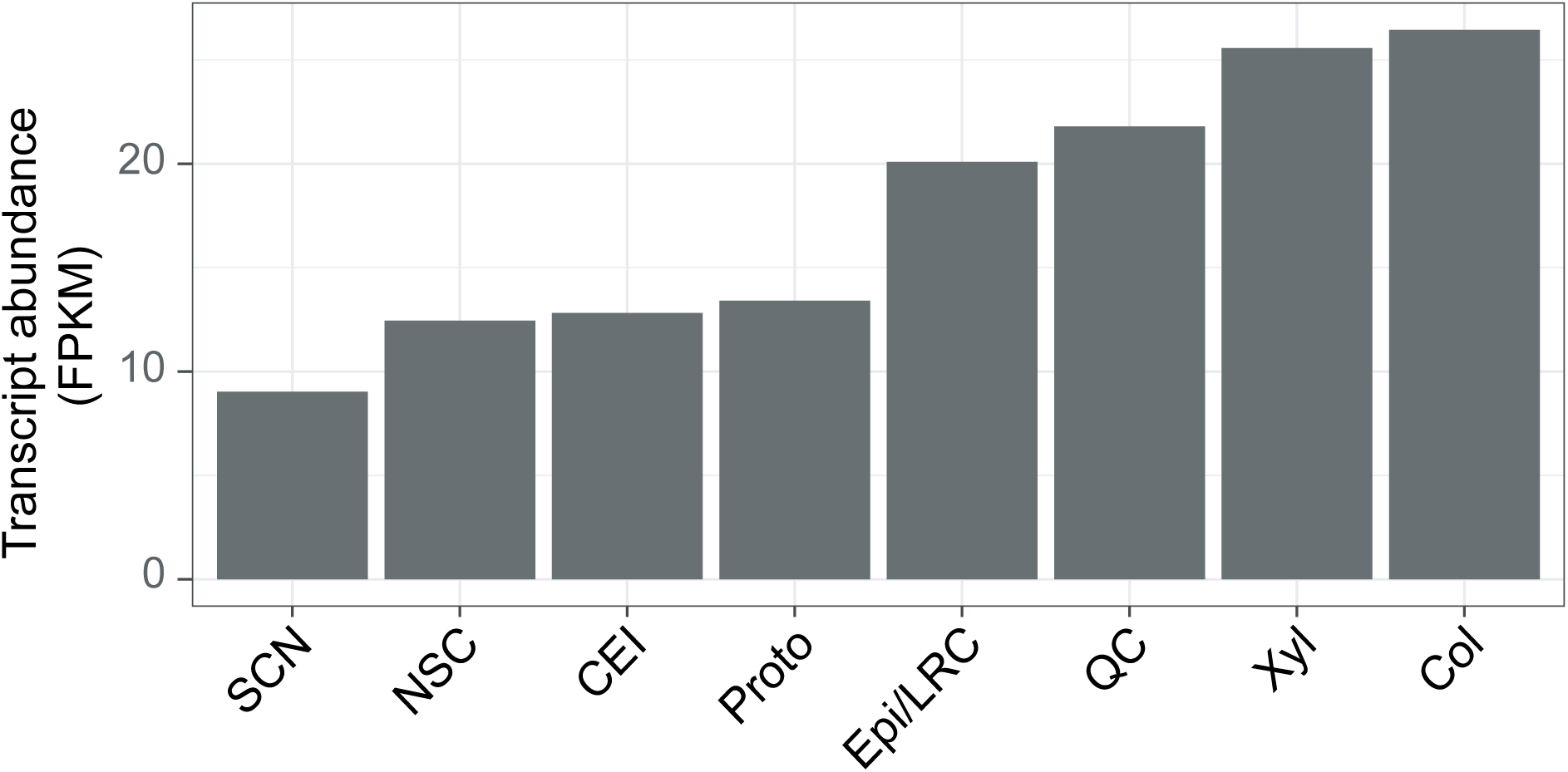
Transcript abundance of GAUT10 in different cell types of *Arabidopsis thaliana* root meristem.

**Figure S2.**
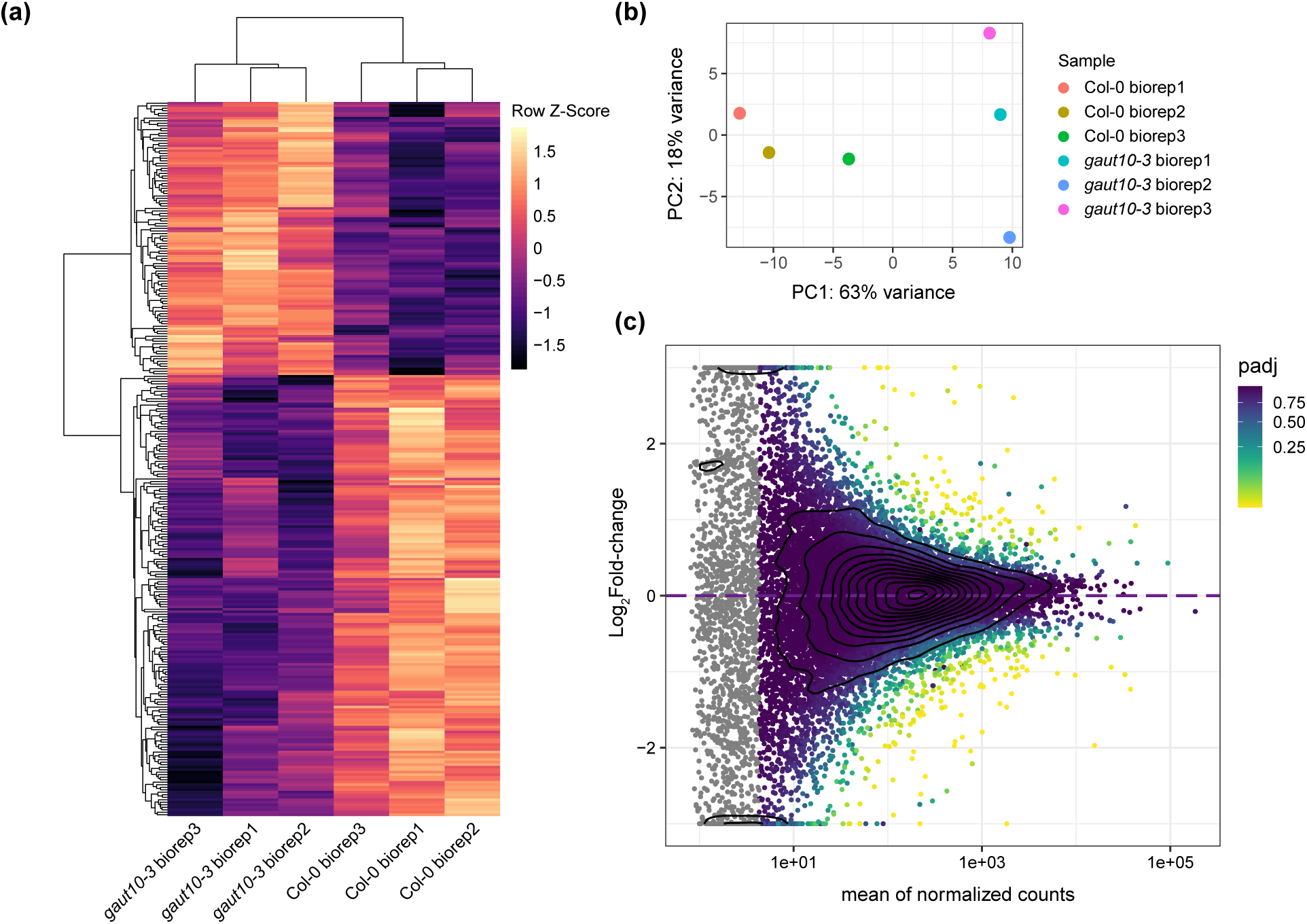
Transcriptomics analysis of 5-days old *gaut10-3* and Col-0 roots across three biological replicates. (a) Heat map shows the relative transcript abundance of all 285 differentially expressed genes clustered based on their row normalized Z-scores and biological replicates. (b) Principal component analysis (PCA) from DESeq2 package shows clustering of three biological replicates across two genotypes in a plane with PC1 as x-axis and PC2 as y-axis. (c) Dispersion plot from DESeq2 analysis showing mean normalized read counts in x-axis and Log_2_FoldChange in y-axis, with genes color coded with their adjusted *P* values, where lighter shaded dots represent statistically significant DEGs.

**Figure S3.**
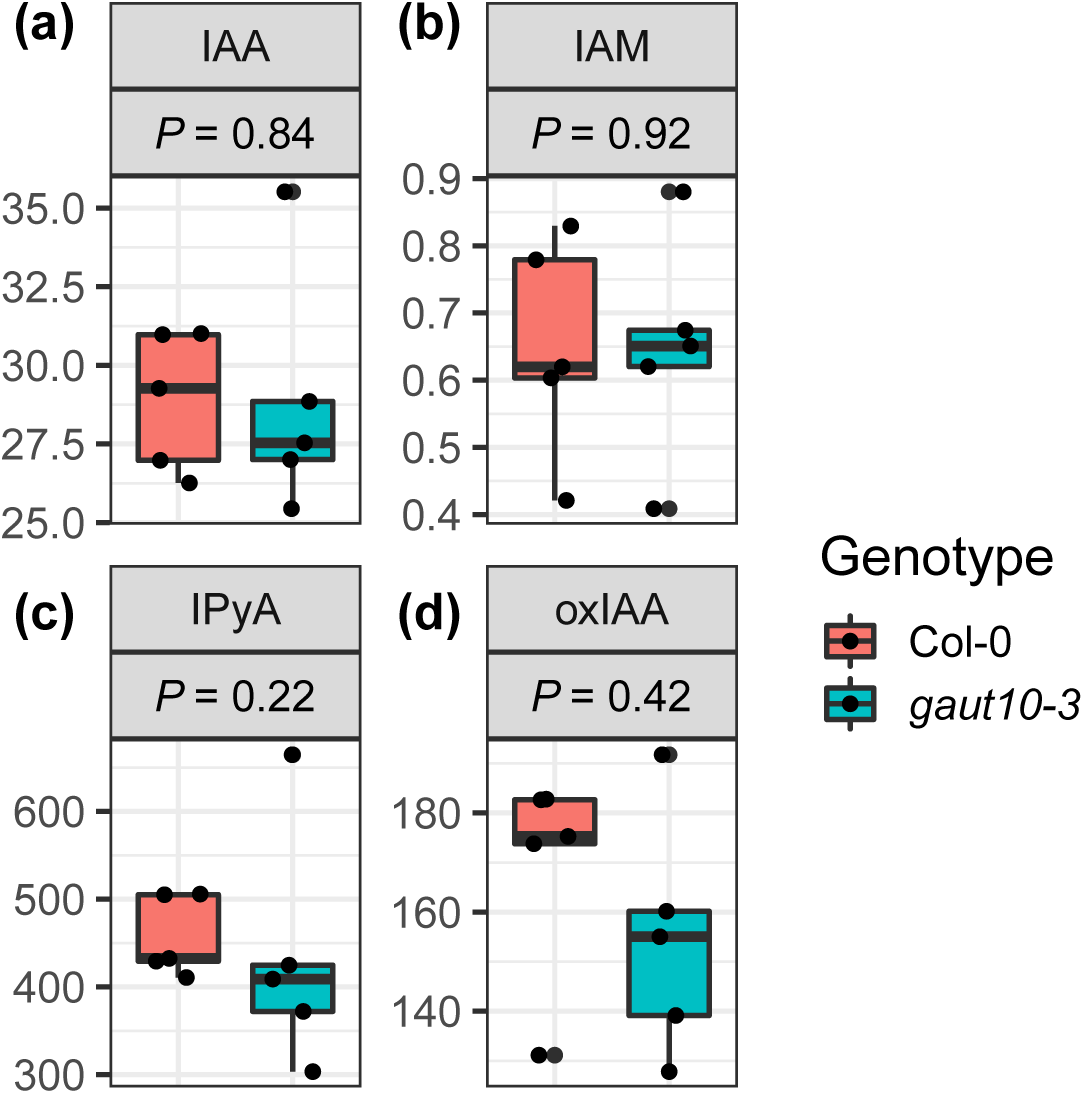
Box-whisker plots show differences in concentrations (in picomoles per gram of fresh weight) of auxin metabolites between five-day-old *gaut10-3* and Col-0 roots averaged across 5 biological replicates. Two sample nonparametric Wilcoxon rank sum tests followed by Benjamini-Hochberg correction for multiple testing are done to identify significantly enriched or reduced metabolites with *P* ≤ 0.1. The tested metabolites with their abbreviations are as follows: indole-3-acetaldoxime acid (IAM), indole-3-pyruvic acid (IPyA), auxin (indole-3-acetic acid, IAA), 2-oxindole-3-acetic acid (oxIAA).

## Notes

### Competing Interest Statement

The authors have declared no competing interest.

